# Requirement for TRanslocon-Associated Protein (TRAP) α in insulin biogenesis

**DOI:** 10.1101/531491

**Authors:** Xin Li, Omar A. Itani, Leena Haataja, Kathleen J. Dumas, Jing Yang, Jeeyeon Cha, Stephane Flibotte, Hung-Jen Shih, Jialu Xu, Ling Qi, Peter Arvan, Ming Liu, Patrick J. Hu

## Abstract

The mechanistic basis for the biogenesis of peptide hormones and growth factors is poorly understood. Here we show that the conserved endoplasmic reticulum (ER) membrane translocon-associated protein (TRAP) α, also known as signal sequence receptor 1 (SSR1)^1^, plays a critical role in the biosynthesis of insulin. A genetic screen in the nematode *Caenorhabditis elegans* revealed *trap-1*, which encodes the *C. elegans* TRAPα ortholog, as a modifier of DAF-2 insulin receptor (InsR) signaling. Genetic analysis indicates that TRAP-1 acts upstream of DAF-2/InsR to control *C. elegans* development. Endogenous *C. elegans* TRAP-1 and mammalian TRAPα both localized to the ER. In pancreatic beta cells, TRAPα deletion impaired preproinsulin translocation but did not affect the synthesis of α_1_-antitrypsin, indicating that TRAPα selectively influences the translocation of a subset of secreted proteins. Surprisingly, loss of TRAPα function also resulted in disruption of distal steps in insulin biogenesis including proinsulin processing and secretion. These results show that TRAPα assists in the ER translocation of preproinsulin and unveil unanticipated additional consequences of TRAPα loss-of-function on the intracellular trafficking and maturation of proinsulin. The association of common intronic single nucleotide variants in the human TRAPα gene with susceptibility to Type 2 diabetes and pancreatic beta cell dysfunction^2^ suggests that impairment of preproinsulin translocation and proinsulin trafficking may contribute to the pathogenesis of Type 2 diabetes.

---

The *C. elegans* DAF-2/InsR pathway prevents dauer diapause through a conserved phosphoinositide 3-kinase (PI3K)/Akt pathway that inhibits the FoxO transcription factor DAF-16. In the context of reduced DAF-2/InsR signaling, cytoplasmic DAF-16/FoxO translocates to the nucleus and promotes dauer arrest through transcriptional regulation^3,4^. In a genetic screen for suppressors of the dauer-constitutive phenotype of an *eak-7;akt-1* double mutant strain that exhibits reduced DAF-2/InsR signaling and increased DAF-16/FoxO activity^5–8^, we isolated a strain containing a ~3.5 kb deletion, *dpDf665*, that spanned the *trap-1* gene as well as three exons of the upstream gene Y71F9AL.1 (Figure 1A). Three independent *trap-1* null mutations (Figure S1) phenocopied the *dpDf665* deletion (Figure 1B), whereas a null mutation in Y71F9AL.1 did not (Figure S2), indicating that the mutant phenotype is a consequence of *trap-1* deletion. *trap-1* mutation suppressed the dauer-constitutive phenotype of mutants with reduced DAF-2/InsR signaling (Figure 1C) but not the phenotype of mutants with reduced signaling in other pathways that inhibit dauer diapause^3,9^ (Figure S3). Furthermore, *trap-1* mutation impaired the induction of DAF-16/FoxO target genes caused by DAF-2/InsR mutation^10^ (Figures 1D-F). Thus, TRAP-1 promotes dauer arrest by specifically antagonizing the DAF-2/InsR pathway.

**Figure 1.**
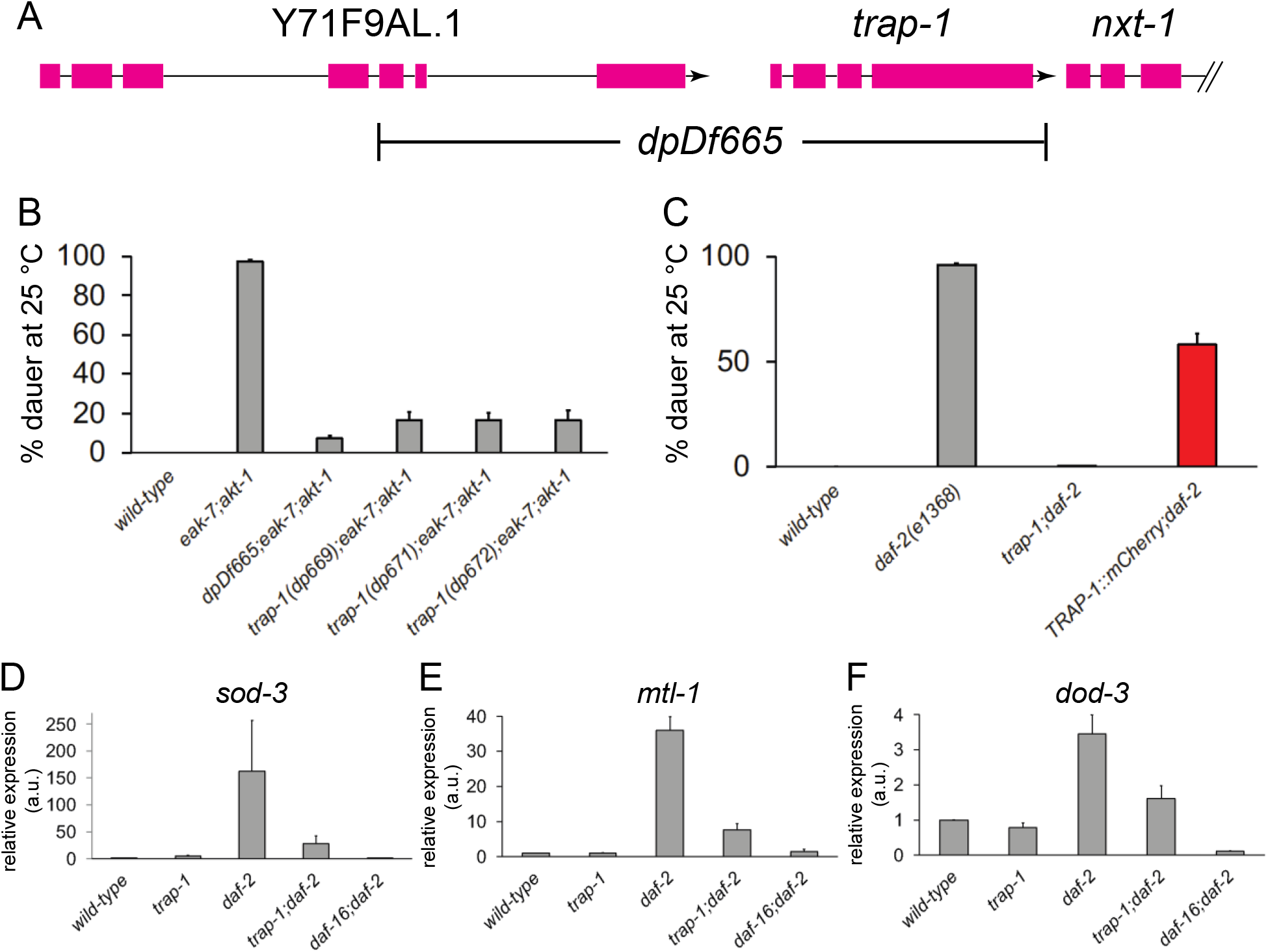
*trap-1* loss-of-function enhances DAF-2/InsR signaling. A. Schematic of the *trap-1* genomic region and the *dpDf665* deletion allele identified in a genetic screen. B. *trap-1* null alleles *dp669, dp671*, and *dp672* (Figure S1) phenocopy dauer suppression caused by *dpDf665* deletion. C. The *trap-1(dp672)* null mutation suppresses the dauer-constitutive phenotype of a *daf-2/InsR* loss-of-function mutant, and a TRAP-1::mCherry fusion protein is functional. D.-F. The *trap-1(dp672)* null mutation inhibits the expression of the DAF-16/FoxO target genes D. *sod-3*, E. *mtl-1*, and F. *dod-3*. a.u., arbitrary units.

TRAP-1 is orthologous to mammalian TRAPα/SSR1 (henceforth referred to as “TRAPα”), a transmembrane ER protein identified based on its interaction with the preprolactin signal peptide during *in vitro* protein translocation^1,11^ (Figure S4). TRAPα physically associates with three other conserved ER transmembrane proteins (TRAPβ, γ, and δ) to form the TRAP complex^12^. A functional single-copy TRAP-1::mCherry fusion protein generated by CRISPR/Cas9-mediated knock-in (Figure 1C) is expressed in several tissues in *C. elegans* embryos, larvae, and adult animals (Figures 2A-C). Coexpression of TRAP-1::mCherry with the ER protein signal peptidase fused to GFP^13^ revealed that endogenous TRAP-1 localizes to the ER (Figure 2D).

**Figure 2.**
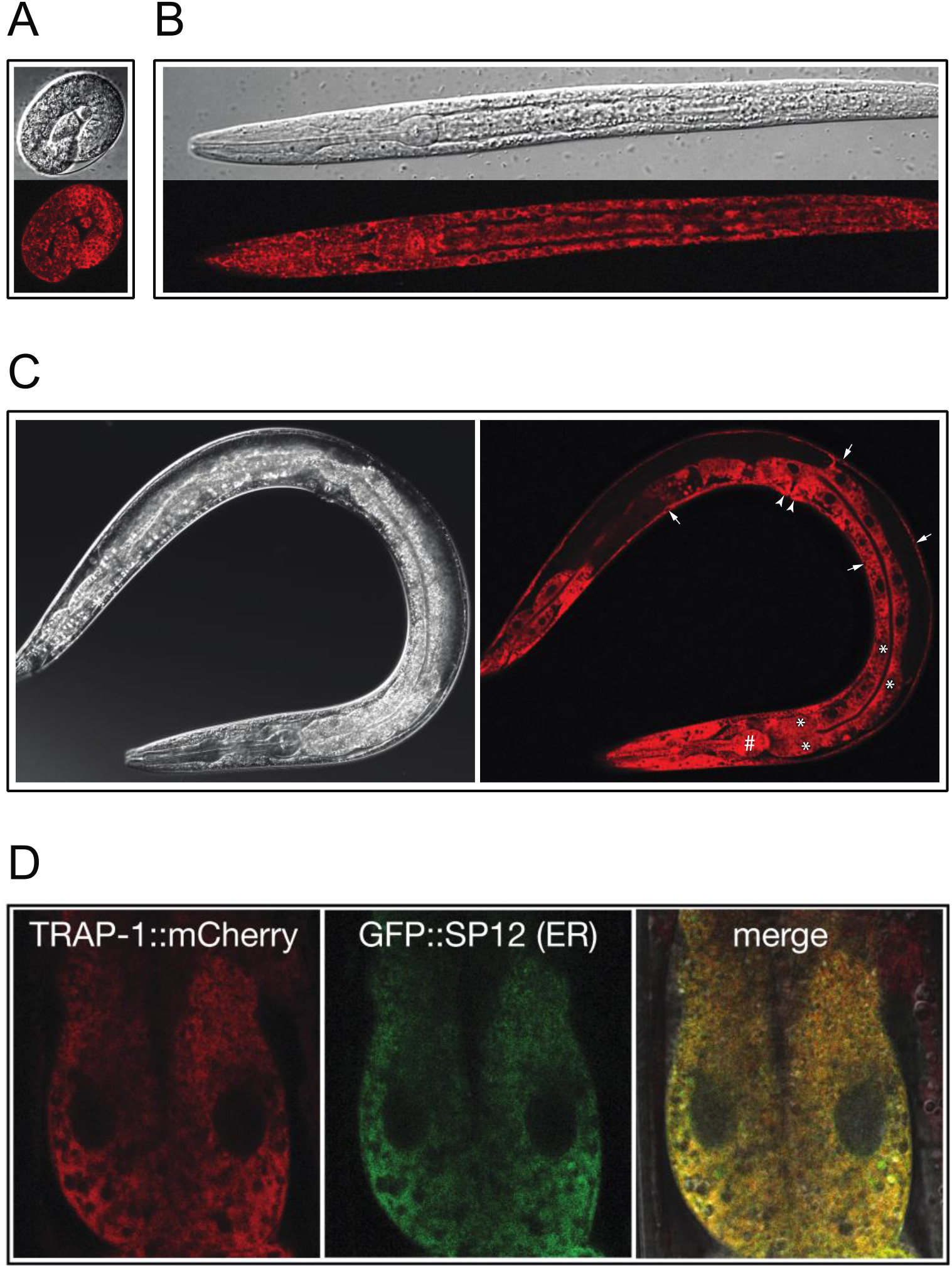
Spatiotemporal expression of a functional TRAP-1::mCherry fusion protein. TRAP-1::mCherry is expressed widely in A. embryos, B. larvae, and C. adult animals. Differential interference contrast (upper panel in A. and B., left panel in C.) and fluorescence (lower panel in A. and B., right panel in C.) images are shown of representative animals. In adults (C.), TRAP-1::mCherry is expressed in the pharynx (hashtag), intestine (asterisks), hypodermis (arrows), and vulva (arrowheads). D. TRAP-1::mCherry colocalizes with the ER signal peptidase GFP-SP12. Two anterior intestinal cells are shown.

Although the biological function of mammalian TRAPα has not been established, it is thought to play a role in cotranslational ER translocation based on its interaction with the preprolactin signal peptide *in vitro*^11^, its necessity for the *in vitro* translocation of prion protein^14^, and its physical proximity to the Sec61 protein translocation channel^15–19^. As the expanded *C. elegans* insulin-like gene family encodes 40 peptides, some of which enhance dauer arrest by antagonizing DAF-2/InsR signaling^20,21^, we hypothesized that TRAP-1 influences DAF-2/InsR signaling by promoting the ER-based biosynthesis of one or more of these insulin-like negative regulators of DAF-2/InsR^20^. If this model is correct, then DAF-2/InsR mutations that impair receptor function downstream of ligand binding should be resistant to suppression by *trap-1* mutation, as these mutant DAF-2/InsR receptors would be refractory to changes in ligand-mediated activity caused by *trap-1* mutation. We therefore tested the effect of *trap-1* mutation on the dauer-constitutive phenotype of eight distinct *daf-2/InsR* loss-of-function alleles, seven of which affect amino acid residues that are conserved in the human InsR^22^ (Table S1).

Whereas *trap-1* mutation suppressed the dauer-constitutive phenotype of the *daf-2* alleles *e1368* and *m212* (Figure S5A and Table S1), both of which encode receptors with missense mutations in the extracellular ligand-binding domain^22,23^, the dauer-constitutive phenotypes of the other six *daf-2* alleles were not affected by *trap-1* mutation. The functional consequences of four of these six DAF-2/InsR mutations can be inferred from data on human InsRs with point mutations affecting the corresponding conserved residues (Table S1). A heterozygous mutation in the human InsR kinase domain corresponding to *daf-2(e1391)*^23^ was found in a patient with autosomal dominant Type A insulin-resistant diabetes mellitus with acanthosis nigricans (IRAN)^24^ and results in a severe reduction in InsR *in vitro* kinase activity and autophosphorylation^25^. Another Type A IRAN patient was found to be homozygous for a missense mutation in the InsR extracellular domain corresponding to *daf-2(m579)*^26^ that reduces the affinity of InsR for insulin by three-to-five fold^27^. A missense mutation affecting the conserved residue in the human InsR corresponding to the glycine mutated in *daf-2(m596)* was identified in a patient with insulin resistance and leprechaunism^28^ and reduces the amount of InsR present at the plasma membrane by more than 90%^29^. Finally, human InsR harboring a missense mutation affecting the conserved cysteine mutated in *daf-2(m577)*^22^ exhibits a 100-fold reduction in levels of mature tetrameric receptor present at the cell surface compared to wild-type human InsR^30^. The failure of *trap-1* mutation to suppress the dauer-constitutive phenotypes of these *daf-2/ĩnsR* alleles (Figure S5A and Table S1) indicates that TRAP-1 acts upstream of DAF-2/InsR and is consistent with a role for TRAP-1 in the biogenesis of insulin-like peptide ligands that antagonize DAF-2/InsR.

To further test the hypothesis that TRAP-1 promotes the biogenesis of antagonistic DAF-2/InsR ligands, we determined the effect of *trap-1* mutation on the dauer-constitutive phenotype of animals with mutations in three genes encoding DAF-2/InsR agonist peptides^21^. *trap-1* mutation partially suppressed the dauer-constitutive phenotype of *ins-4 ins-6;daf-28* triple mutants (Figure S5B), consistent with a model whereby TRAP-1 promotes antagonist insulin-like peptide ligand action.

As human insulin antagonizes DAF-2/InsR signaling when it is expressed in *C. elegans*^20^, we explored the possibility that mammalian TRAPα promotes insulin biogenesis. Endogenous TRAPα colocalized with KDEL ER resident proteins but not with the Golgi marker GM130^31^ in rat INS 832/13 pancreatic beta cells (Figure 3A). To assess the role of TRAPα in preproinsulin translocation, we generated INS 832/13 cells lacking TRAPα using CRISPR/Cas9-based genome editing. Normally, preproinsulin is nearly undetectable by immunoblotting with anti-proinsulin antibodies (Figure 3B, first two lanes), but TRAPα knockout (KO) resulted in detectable amounts of preproinsulin (Figure 3B, third lane) that increased further upon pretreatment of cells with the proteasome inhibitor MG132 (Figure 3B, last lane).

**Figure 3.**
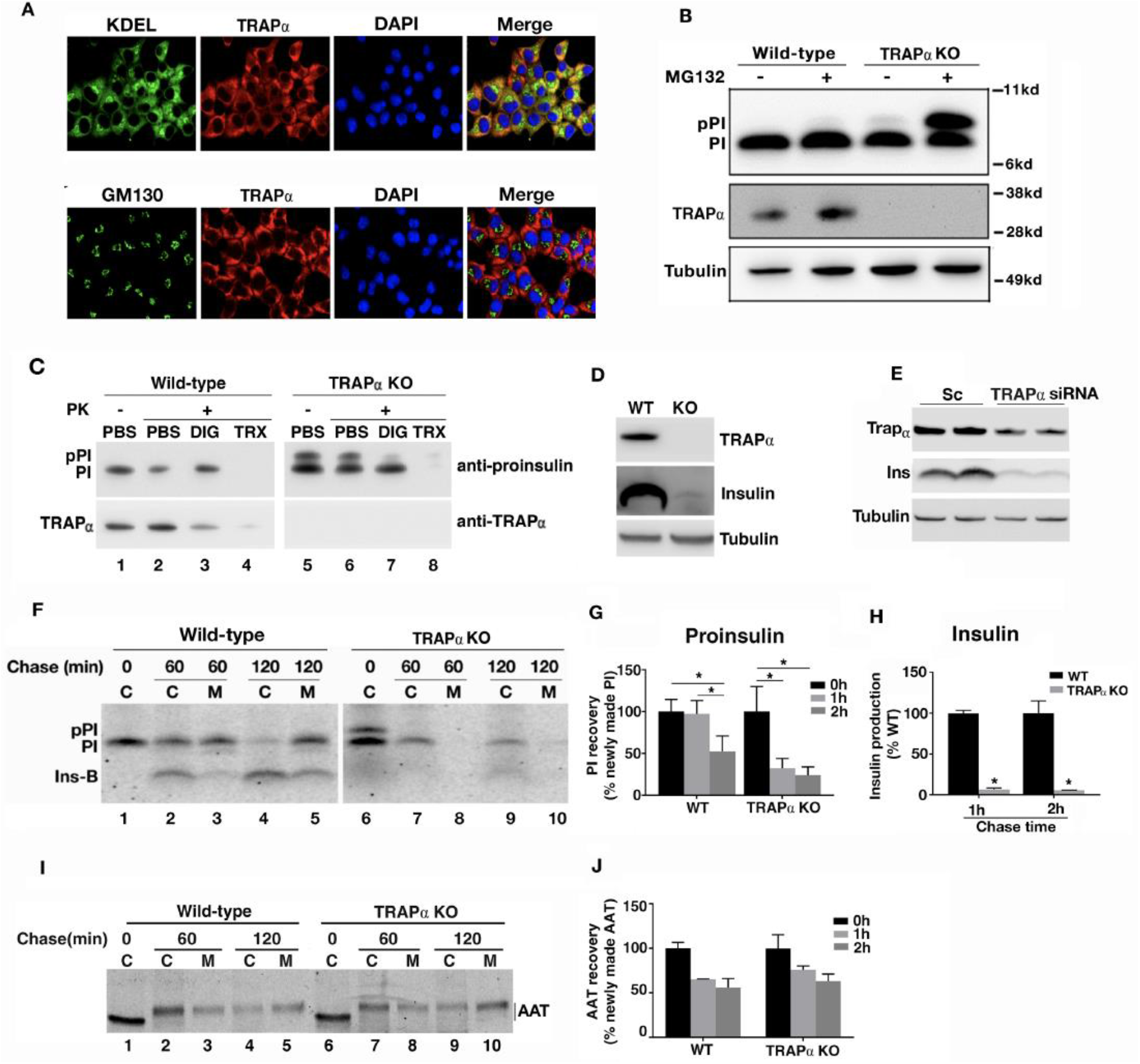
TRAPα promotes preproinsulin ER translocation, insulin biogenesis, and insulin secretion in INS 832/13 cells. A. Immunostaining of INS 832/13 cells with anti-TRAPα antibodies reveals colocalization with ER proteins recognized by anti-KDEL antibodies (upper panels) but not with the Golgi protein GM130 (lower panels). Nuclei are stained with DAPI. B. SDS-PAGE and anti-proinsulin immunoblotting of lysates from INS 832/13 wild-type or TRAPα KO cells. Cells were untreated or treated with the proteasome inhibitor MG132 as indicated. Abbreviations: pPI, preproinsulin; PI, proinsulin. C. SDS-PAGE and anti-proinsulin immunoblotting of lysates from INS 832/13 wild-type or TRAPα KO cells after pretreatment with phosphate-buffered saline (PBS), digitonin (DIG), or Triton X-100 (TRX) and subsequent exposure to Proteinase K (PK). D. SDS-PAGE and anti-insulin immunoblotting of lysates from INS 832/13 wild-type and TRAPα KO cells. E. SDS-PAGE and anti-insulin immunoblotting of lysates from INS 832/13 wild-type cells that were transfected with either scrambled (Sc) or TRAPα siRNAs. Duplicate samples were run. F. SDS-PAGE of anti-insulin immunoprecipitates of cell lysates (C) or conditioned media (M) from INS 832/13 wild-type or TRAPα KO cells after pulse-labeling with ^35^S-Met/Cys and chase for the indicated times. Abbreviations: pPI, preproinsulin; PI, proinsulin; Ins-B, insulin. Quantification of total proinsulin and insulin (C + M) is shown in G. and H., respectively. I. SDS-PAGE of anti-α1-antitrypsin (AAT) immunoprecipitates of cell lysates (C) or conditioned media (M) from INS 832/13 wild-type or TRAPα KO cells after pulse-labeling and chase as described for F. INS 832/13 wild-type or TRAPα KO cells were transfected with a plasmid encoding AAT 48 hours prior to pulse-labeling. J. Quantification of total AAT (C + M) at the indicated time points.

These data could be consistent with a requirement for TRAPα in the translocation of preproinsulin such that in the absence of TRAPα, untranslocated preproinsulin is degraded by the proteasome. Alternatively, the data in Figure 3B could be consistent with impaired preproinsulin processing by signal peptidase in the ER. To distinguish between a translocation defect and a signal peptidase processing defect in TRAPα KO cells, we determined the protease sensitivity of preproinsulin in TRAPα KO cells after selective permeabilization of the plasma membrane (Figure 3C). In wild-type cells, preproinsulin was fully translocated into the ER and processed to proinsulin that was protease-resistant after plasma membrane permeabilization with digitonin but degraded after full solubilization of internal membranes with Triton X-100 (Figure 3C, third and fourth lanes). In TRAPα KO cells, accumulated preproinsulin was protease-sensitive both after partial as well as complete membrane permeabilization (Figure 3C, seventh and eighth lanes). These data indicate that in TRAPα KO cells, preproinsulin remains untranslocated in the cytosol.

While the steady-state level of proinsulin was not affected by TRAPα KO (Figures 3B and 3C), anti-insulin immunoblotting revealed a surprisingly dramatic reduction in total insulin content in TRAPα KO cells compared to control cells (Figure 3D). TRAPα knockdown by RNAi recapitulated this decrease in steady-state insulin levels (Figure 3E), indicating that this phenotype does not represent an off-target effect of genome editing.

To gain further insight into the role of TRAPα in early events governing insulin biogenesis, we pulse-labeled cells with ^35^S-labeled methionine and cysteine (^35^S-Met/Cys) and analyzed insulin biosynthetic intermediates by immunoprecipitation immediately after labeling or after a chase of 60 or 120 minutes (Figures 3F-H). In wild-type cells, proinsulin (PI) was efficiently labeled with ^35^S-Met/Cys (Figure 3F, lane 1) and processed into mature insulin (Ins-B; lanes 2 and 4). Additionally, both proinsulin and insulin were secreted into the media (M; lanes 3 and 5). Notably, in wild-type cells, preproinsulin (pPI) was not detected, consistent with its rapid translocation and conversion to proinsulin in the ER. In contrast, in TRAPα KO cells, newly synthesized preproinsulin was detectable immediately after labeling (lane 6) but was lost thereafter. While proinsulin was still synthesized (lane 6), its recovery at 60 and 120 minutes after labeling was diminished (compare lanes 7 and 9 to lane 6), and proinsulin was neither secreted intact nor processed to mature insulin intracellularly (lanes 7-10). Therefore, in addition to impairing preproinsulin ER translocation, TRAPα KO also results in reduced proinsulin stability, with dramatically decreased insulin biogenesis and secretion.

To elucidate the contribution of the proteasome to increased proinsulin turnover in TRAPα KO cells, we treated ^35^S-Met/Cys-labeled cells with the proteasome inhibitor MG132. MG132 treatment increased intracellular proinsulin levels (Figure S6, compare lane 13 to lane 11 and lane 17 to lane 15). However, this increase did not rescue defects in proinsulin secretion and processing to mature insulin. Therefore, TRAPα KO impairs the intracellular trafficking of proinsulin and the biogenesis of insulin independent of its effect on proinsulin turnover.

To determine whether the proinsulin trafficking defect observed in TRAPα KO cells was due to a general defect in the secretory pathway, we analyzed the biogenesis and secretion of α1-antitrypsin (AAT) in wild-type and TRAPα KO cells. TRAPα KO affected neither the biogenesis nor the secretion of AAT (Figures 3I-J). These findings indicate that the secretory pathway is functionally intact in TRAPα KO cells and also demonstrate that TRAPα promotes the ER translocation and subsequent trafficking of only a subset of secreted proteins.

We also used immunofluorescence to independently examine the consequences of TRAPα KO on insulin biogenesis. In wild-type cells, insulin (Figure 4A) and proinsulin (Figure 4B) were readily detectable. However, TRAPα KO resulted in a dramatic decrease in insulin content (Figure 4A), although proinsulin was still detectable (Figure 4B).

**Figure 4.**
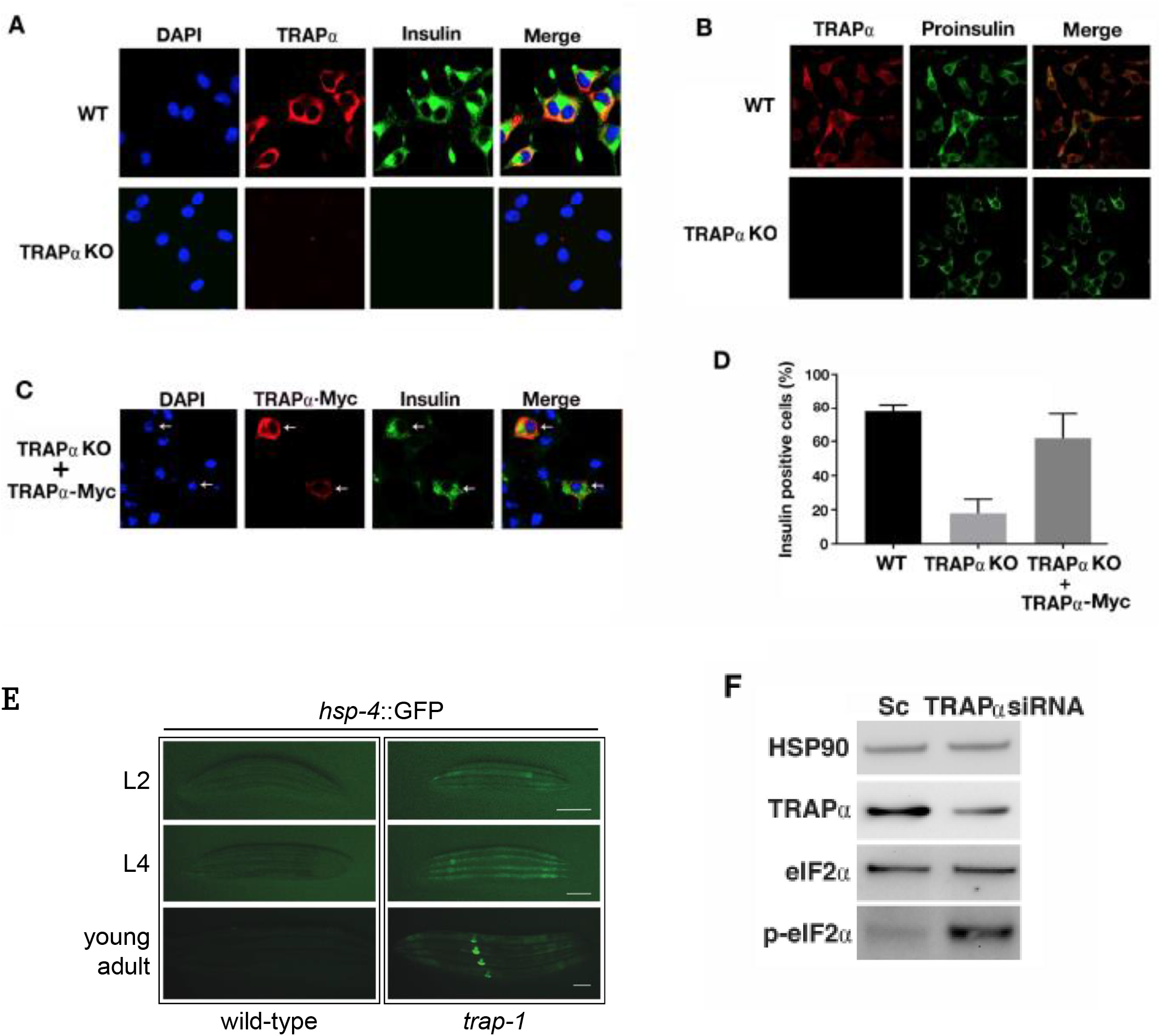
TRAPα promotes insulin biosynthesis and reduces ER stress. A. Immunostaining of INS832/13 wild-type (WT) or TRAPα KO cells with anti-TRAPα and A. anti-insulin or B. anti-proinsulin antibodies. Nuclei are stained with DAPI. C. Immunostaining of TRAPα KO cells with anti-Myc and anti-insulin antibodies after transfection with a plasmid encoding a Myc-tagged TRAPα cDNA. Nuclei are stained with DAPI. D. Quantification of insulin-positive cells in INS832/13 wild-type, TRAPα KO, and TRAPα KO cells expressing exogenous Myc-tagged TRAPα. E. Expression of the ER stress reporter *hsp-4::GFP* in second (L2) and fourth stage (L4) larvae and young adult wild-type (left panels) and *trap-1(dp672)*null mutant (right panels) animals. At each stage, images of wild-type and *trap-1* mutant animals were captured with equivalent exposure times. Scale bar: 100 microns. F. SDS-PAGE and anti-phospho-Ser51-eIF2 immunoblotting of lysates from INS 832/13 wild-type cells that were transfected with either scrambled (Sc) or TRAPα siRNA as indicated.

Reintroducing epitope-tagged TRAPα into TRAPα KO cells rescued insulin production in transfected cells (Figures 4C-D), and infection of TRAPα KO cells with adenovirus encoding TRAPα at increasing multiplicities of infection revealed that this rescue was dose-dependent (Figure S7).

Our finding that TRAPα activity is required for efficient insulin biogenesis and secretion was unexpected. Defects in proinsulin processing and secretion could be an indirect consequence of TRAPα dysfunction in the ER. In mouse fibroblasts, TRAPα physically associates with misfolded ER-associated degradation (ERAD) substrates^32^, suggesting that TRAPα may play a role in reducing ER stress. Thus, in pancreatic beta cells, loss of TRAPα may cause abnormalities in proinsulin processing and secretion (Figures 3F-H) due to a general increase in ER stress.

To determine whether *C. elegans* TRAP-1 and mammalian TRAPα play roles in mitigating ER stress, we assayed for constitutive activation of the unfolded protein response in animals and beta cells lacking TRAP-1 or TRAPα, respectively. In *C. elegans*, transcription of *hsp-4*, which encodes a homolog of the human ER chaperone BiP, is induced by ER stress^33^. Indeed, while green fluorescence was not detectable in wild-type *hsp-4::*GFP reporter animals grown under normal culture conditions, it was readily visualized in *trap-1* mutant animals harboring the same transgene (Figure 4E). Similarly, increased phosphorylation of eIF2α at serine 51, which is phosphorylated by PERK in response to ER stress^34^, was detected in INS 832/13 cells with reduced TRAPα expression but not in wild-type INS 832/13 cells (Figure 4F). Thus, we cannot exclude the possibility that TRAPα may indirectly influence distal events in insulin biogenesis through effects on ER stress that are not similarly consequential for α1-antitrypsin (Figures 3I-J).

Although the role of ER translocation in governing insulin biogenesis has been underappreciated, hints of its importance have emerged from studies on rare missense mutations in the signal peptide of human preproinsulin that are associated with congenital diabetes^35^. The current work provides insight into the importance of the ER translocation machinery in insulin biogenesis by establishing preproinsulin as the first bonafide client protein for TRAPα. Our data are consistent with a model whereby TRAPα directly promotes preproinsulin translocation and indirectly influences proinsulin maturation and insulin secretion, possibly through its effects on ER stress. Variation in TRAPα expression and/or activity may be generally relevant to the pathogenesis of Type 2 diabetes, as several common intronic single nucleotide variants in the human *Ssr1* gene (encoding TRAPα) are associated with Type 2 diabetes risk and pancreatic beta cell dysfunction^2^. Intriguingly, genes encoding TRAP subunits are among the most highly upregulated genes in pancreatic beta cells exposed to high glucose concentrations^36^, and the transcriptional induction of TRAPα, ß, and γ in pancreatic beta cells requires the unfolded protein response transducer IRE1α^37^. Thus, upregulation of TRAPα in pancreatic beta cells in the context of hyperglycemia may promote metabolic stability by both enhancing insulin biogenesis and attenuating ER stress. Understanding the mechanistic basis for TRAPα action in insulin biogenesis may lead to new approaches to prevent and treat Type 2 diabetes.

As mammalian TRAPα is expressed widely^38^, it surely must have biological functions that are distinct from its role in insulin biogenesis. These may include promoting the translocation of other proteins that enter the secretory pathway (such as PrP^14^) and/or participating in ERAD^32^. Mice homozygous for a C-terminal TRAPα-*ShBle-LacZ* fusion allele die perinatally, and homozygous TRAPα-*ShBle-LacZ* embryos are small and have cardiac outflow tract and endocardial cushion defects^38^. Therefore, TRAPα likely plays a role in the translocation of other secreted proteins that play important roles in embryonic development. Defining specific features of translocated proteins that confer TRAPα dependence^14,39^ will facilitate the identification of additional TRAPα client proteins and the elucidation of other biological functions of TRAPα.

**Supplementary Information** is linked to the online version of the paper.

## Acknowledgements

This work was supported by American Heart Association Postdoctoral Fellowship Award 14POST20390031 (O.A.I.); NIH R01DK120047 (L.Q.), R01DK111174 (L.Q., P.A., and M.L.), R01DK048280 (P.A.), and R01DK088856 (M.L.); National Natural Science Foundation of China grants 81070629, 81570699, 81620108004, and 81370895 (M.L.); and NIH R01AG041177, American Cancer Society Research Scholar Grant 119640-RSG-10-132-01-DDC, and American Heart Association Innovative Research Grant 11IRG5170009 (P.J.H.). We acknowledge support from the Michigan Diabetes Research Center Morphology Core (NIH P30 DK020572) and thank Roland Stein for comments on the manuscript.

## Author contributions

Conceived of the project: O.A.I., P.A., M.L., P.J.H.; performed experiments: X.L., O.A.I., L.H., K.J.D., J.Y., J.C., H.-J.S., J.X.; analyzed data: X.L., O.A.I., L.H., K.J.D., J.C., S.F., H.-J.S., L.Q., P.A., M.L., P.J.H.; wrote the manuscript: P.A., M.L., P.J.H.

## Competing interests

The authors declare no competing interests.

## Materials and correspondence

Requests for materials and other correspondence should be addressed to Patrick J. Hu.

## METHODS

### Generation and maintenance of *C. elegans* strains

Strains were maintained on standard nematode growth media (NGM) plates seeded with *Escherichia coli* OP50. Double and triple mutant strains were constructed using conventional methods. *trap-1* mutant alleles were generated using CRISPR/Cas9-based genome editing^1^. Two guide RNAs complementary to sequences in exon 2 were used. Molecular analysis of *trap-1* mutants is described in Figure S1. DNA encoding a mCherry epitope tag was inserted in-frame at the 3’ end of the *trap-1* open reading frame using CRISPR/Cas9-based homology-directed genome editing as described^2^. Red fluorescent animals were isolated, and the mCherry insertion was verified by Sanger sequencing of PCR products spanning the insertion site.

### Genetic screen for modifiers of *C. elegans* DAF-2/InsR signaling

The suppressor of *eak-7;akt-1 (seak)* screen was performed by mutagenizing *eak-7;akt-1* double mutants with *N*-ethyl-*N*-nitrosourea (ENU) and screening for rare animals in the F2 generation that did not arrest as dauers as described^3^. Genomic DNA isolated from *eak-7;akt-1* suppressor strains was sequenced and analyzed as described^3^.

### Dauer arrest assays

Dauer arrest assays were performed at 25°C as previously described^4^.

### Real-time quantitative PCR (qPCR)

qPCR was performed as previously described^5^.

### *C. elegans* fluorescence microscopy

Animals were mounted on slides layered with a thin 3% agarose pad containing 25 mM sodium azide. Images were captured on a Leica inverted SP5 × inverted confocal microscope using LAS AF software (Leica).

### Reagents

Rat monoclonal anti-KDEL and rabbit monoclonal anti-GM130 antibodies were from Abcam (ab50601 and ab52649, respectively). Rabbit polyclonal anti-TRAPα antibodies were from Novus Biologicals (NBP1-86912). A mouse monoclonal antibody that recognizes a human proinsulin C-peptide—A-chain junction peptide (GSLQKRGIVE) was raised by Abmart. Mouse monoclonal anti-tubulin antibodies were from Sigma (T5168). Guinea pig polyclonal anti-porcine insulin antibodies were from Millipore. Rabbit polyclonal anti-α1-antitrypsin (AAT) antibodies were from Dako (A001202-2). Rabbit polyclonal anti-HSP90 (#4874S), anti-eIF2 (#9722S), and anti-phospho-Ser51 eIF2 antibodies (#9721) were from Cell Signaling. ^35^S-Met/Cys was from ICN. Dithiothreitol (DTT), protein A-agarose, digitonin, N-ethylmaleimide (NEM), and other chemical reagents were from Sigma-Aldrich. 4-12% gradient SDS-polyacrylamide gel electrophoresis (SDS-PAGE) was performed using NuPage gels from Thermo Fisher. Met/Cys-deficient Dulbecco’s modified Eagle’s medium (DMEM) and all other tissue culture reagents were from Invitrogen. A cDNA encoding Myc epitope-tagged human TRAPα was synthesized by Integrated DNA Technologies (IDT) and was subcloned into the pTarget mammalian expressing vector (Promega).

### Manipulating TRAPα expression in INS832/13 cells

INS832/13 cells lacking TRAPα were generated using CRISPR/Cas9-mediated genome editing^6^. Single guide RNAs (sgRNAs) were designed using guide design resources available on the Zhang Lab web site (https://zlab.bio/guide-design-resources). Oligonucleotides corresponding to 5’ - CACCGTCTGCACTCGGTGAAGCTTC - 3’ 20-nt guide sequences were annealed and ligated into pSpCas9(BB)-2A-Puro (PX462) v2.0 vector as described^6^. This plasmid was transfected into INS832/13 cells using Lipofectamine 2000 (Thermo Fisher). 48 hours after transfection, cells were plated and cultured with RPMI growth medium containing 1 μg/ml puromycin. Puromycin-resistant clones were screened for loss of TRAPα (“TRAPα KO”) by immunoblotting with anti-TRAPα antibodies. To re-express TRAPα, TRAPα KO cells were transiently transfected with a plasmid encoding Myc-tagged human TRAPα. 48 hours after transfection, insulin content was assayed by immunofluorescence. To reduce TRAPα expression by RNAi, INS832/13 cells were transfected with 20nM TRAPα targeting siRNA (Assay ID 219062, Thermo Fisher) or Negative Control siRNA (AM4611, Thermo Fisher) using Lipofectamine RNAiMAX (Invitrogen). 48 hours after transfection, knockdown efficiency was evaluated by immunoblotting using anti-TRAPα antibodies.

### Cell culture

Cells were plated into 6-well plates one day before transfection. A total of 2 μg plasmid DNA was transfected using Lipofectamine 2000 (Invitrogen). For metabolic labeling, cells were pulse-labeled with ^35^S-Met/Cys for 15 minutes, washed, and chased with label-free media for 0, 60, or 120 minutes as indicated. Immediately thereafter, media and cell lysate were immunoprecipitated with anti-insulin or anti-AAT antibodies, and immunoprecipitates were subjected to 4-12% gradient SDS-PAGE under reducing conditions. Preproinsulin, proinsulin, insulin, and AAT were analyzed using a Typhoon Phosphorimager (GE Healthcare). Band intensities were quantified using NIH ImageJ. For selective plasma membrane permeabilization and Proteinase K digestion, cells were incubated on ice with phosphate-buffered saline (PBS), with or without 10 μg/ml Proteinase K, 0.01% digitonin, or 1% Triton X-100 as indicated for 30 minutes. Cells were harvested, boiled in SDS sample buffer, and analyzed by immunoblotting with anti-proinsulin antibodies.

### Immunofluorescence

Cells were permeabilized with 0.5% Triton X-100, fixed with formaldehyde, blocked, and incubated with anti-TRAPα, anti-KDEL, anti-GM130, antiinsulin, anti-proinsulin, or anti-Myc antibodies. After incubation with fluorophore-conjugated secondary antibodies, specimens were imaged with an Olympus FV500 confocal microscope.

### Immunoblotting

Cell lysates were boiled in sample buffer containing 100 mM DTT for five minutes, resolved by 4–12% gradient SDS-PAGE, and transferred to nitrocellulose. Primary antibodies were used at 1:1000 dilution in TBST (0.1% Tween 20) plus 5% bovine serum albumin, with the exception of anti-HSP90, which was used at 1:2000 dilution. Secondary antibodies [goat anti-rabbit IgG HRP, goat anti-mouse IgG HRP (both from Biorad), and goat anti-guinea pig IgG HRP (Jackson ImmunoResearch)] were used at 1:5000 dilution. Imaging was performed after incubation with Clarity Western ECL Substrate (Bio-Rad #1705061) according to the manufacturer’s instructions.

### Statistical analyses

Statistical analyses were carried out by Student’s t-test or one-way ANOVA using GraphPad Prism 7.

### *hsp-4p.:GFP* reporter assay

Wild-type and *trap-1* mutant *C. elegans* harboring an integrated *hsp-4p.:GFP* transgene^7^ were harvested, incubated in alkaline hypochlorite [500 mM NaOH and 1.2% (v/v) hypochlorite], and vortexed to isolate eggs. Eggs were washed two times with M9 buffer solution and incubated at 20°C on standard NGM plates seeded with *E. coli* OP50. Animals were imaged at L2 larval, L4 larval, and young adult stages on a Leica SP5 × confocal microscope equipped with LAS AF software. Three experimental replicates yielded similar results.

### Data availability

The data that support the findings of this study are available from the corresponding author upon reasonable request.

