## Supplementary material for "Requirement for TRanslocon-Associated Protein (TRAP) α in insulin biogenesis": Figures S1-S7 and Table S1

**EXTENDED DATA**

*trap-1* wild-type exon 2 (guide RNA sequences underlined):

CCGCCGACGTCGTTGACGGAGAAGTCACAGATGACGCGCCAAAGAACTCGCAAGAAGACGATGATCTCACCATCGGAGCCTCCCCAGATGCCGGACTTGCATTCCACTTTGTTCAACCATCTGATGCCAATGTTGTTCGCG

Wild-type protein:

22 …VVDGEVTDDAPKNSQEDDDLTIGASPDAGLAFHFVQPSDANVVR… 65

*dp669* exon 2: two-nucleotide transversion (red); six- and eight-nucleotide deletions (=) resulting in frameshift and premature translation termination (**bold**):

CATC======CGTTGACGGAGAAGTCACAGATGACGCGCCAAAGAACTCGCAAGAAGACGATG========ATCGGAGCCTCCCCAGATGCCGGACTTGCATTCCACTTTGTTCAACCATC**TGA**TGCCAATGTTGTTCGCG

Predicted protein (wild-type sequences in black):

22 …SVDGEVTDDAPKNSQEDDDRSLPRCRTCIPLCSTI*

*dp671* exon 2: 20-nucleotide deletion resulting in frameshift and premature translation termination:

CCGCCG====================ACAGA**TGA**CGCGCCAAAGAACTCGCAAGAAGACGATGATCTCACCATCGGAGCCTCCCCAGATGCCGGACTTGCATTCCACTTTGTTCAACCATCTGATGCCAATGTTGTTCGCG

Predicted protein:

22 …DR*

*dp672* exon 2: three- and eight-nucleotide deletions resulting in frameshift and premature translation termination:

CCGCCGA===CGTTGACGGAGAAGTCACAGATGACGCGCCAAAGAACTCGCAAGAAGACGATG========ATCGGAGCCTCCCCAGATGCCGGACTTGCATTCCACTTTGTTCAACCATC**TGA**TGCCAATGTTGTTCGCG

Predicted protein:

22 …DVDGEVTDDAPKNSQEDDDRSLPRCRTCIPLCSTI*

**Figure S1.** Molecular analysis of the three putative *trap-1* null alleles *dp669, dp671,* and *dp672,* based on sequencing of PCR products spanning exon 2 of the *trap-1* gene.

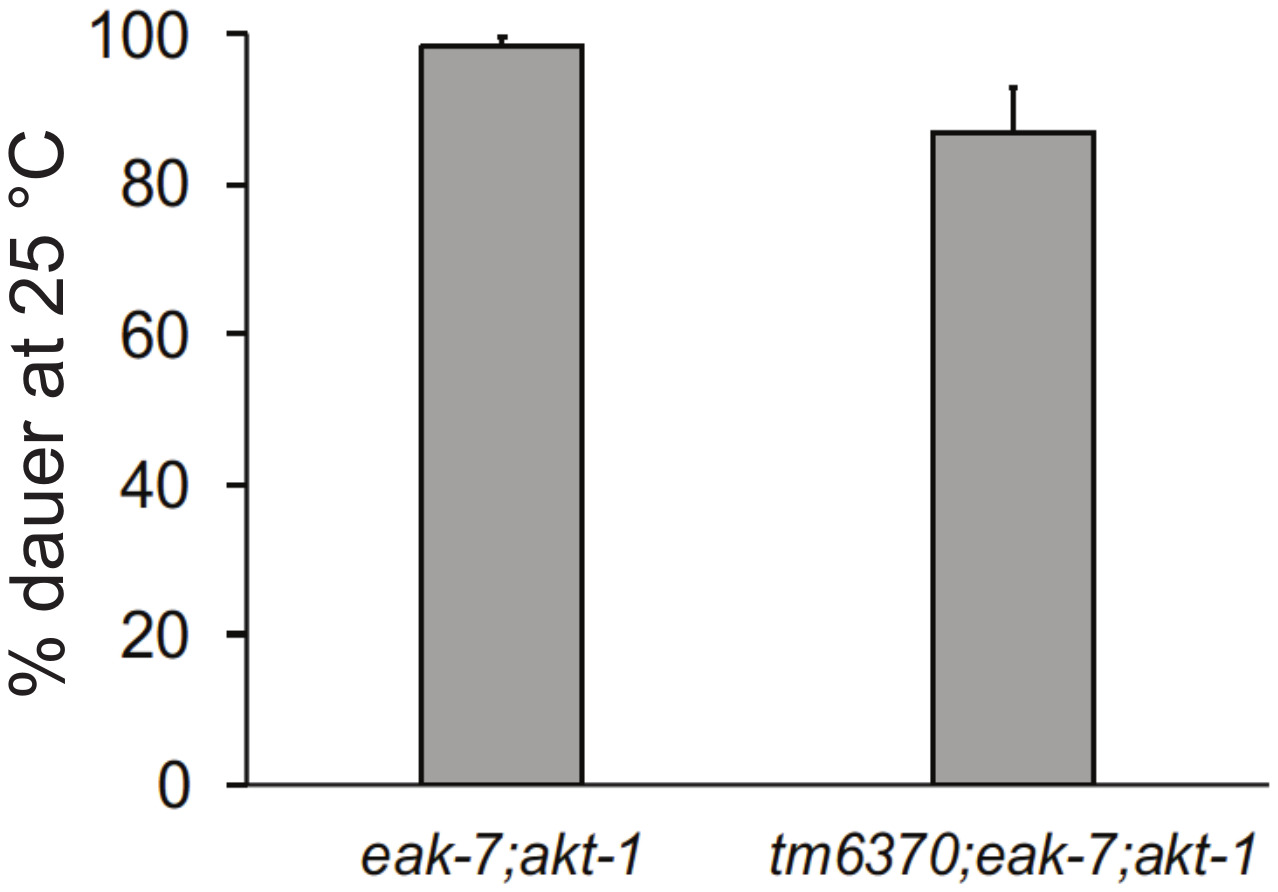

**Figure S2.** The *Y71F9AL.1(tm6370)* deletion allele does not suppress the dauer-constitutive phenotype of *eak-7;akt-1* double mutants (p = 0.08 by Student’s t-test). Error bars: s.e.m.

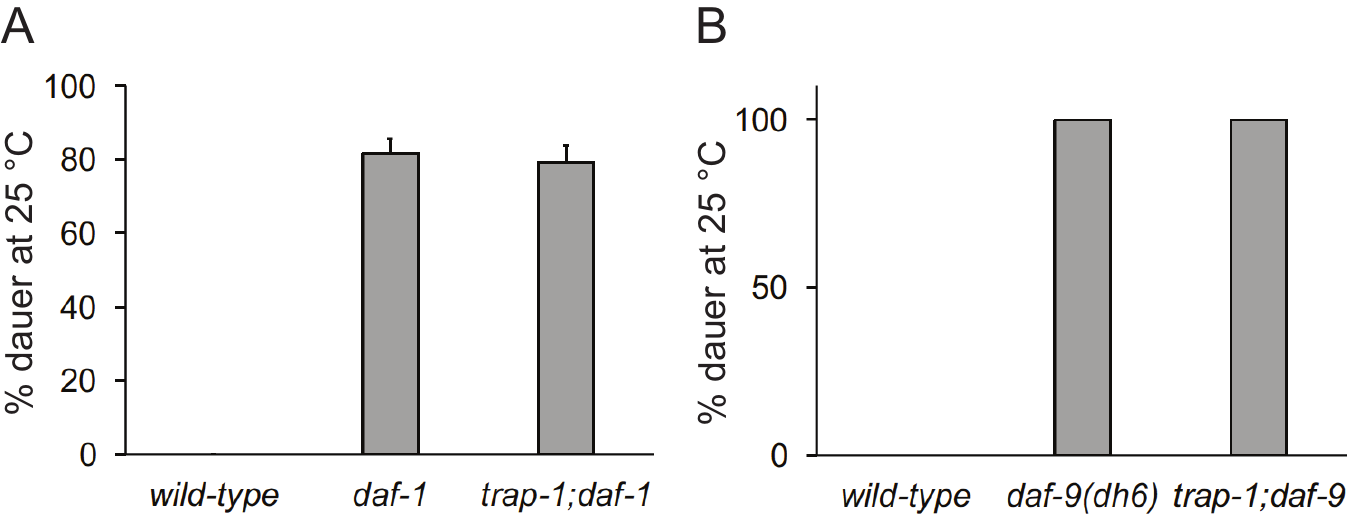

**Figure S3.** A *trap-1* null mutation does not suppress the dauer-constitutive phenotype of A. *daf-1/TGFBRI* or B. *daf-9* null mutants. Error bars: s.e.m.

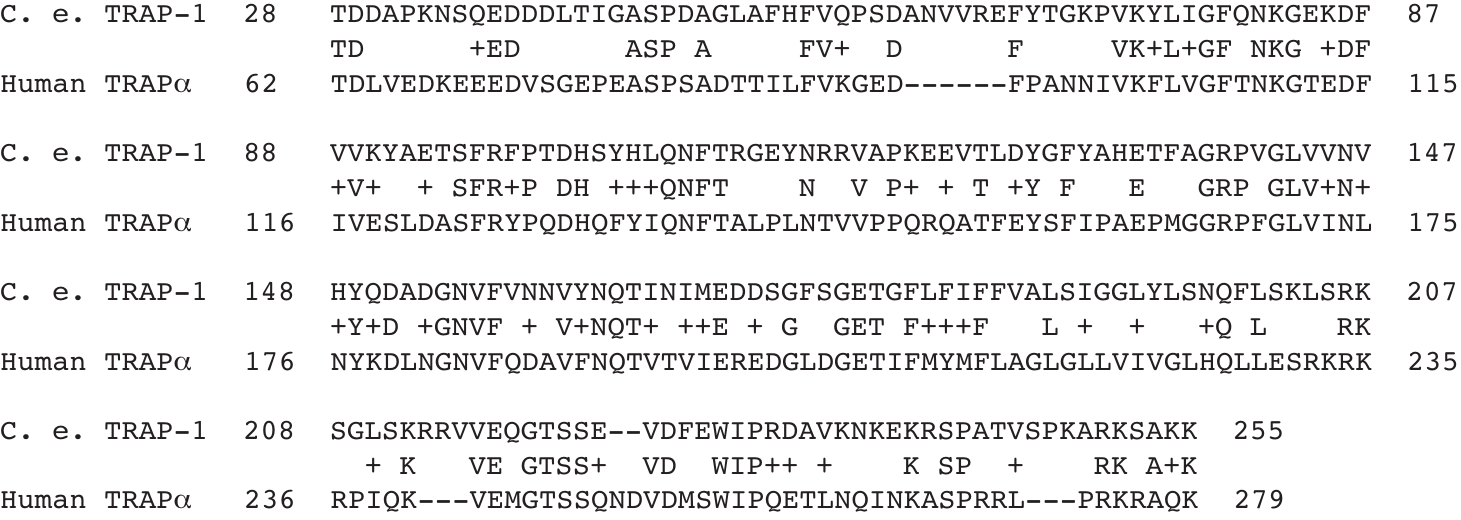

**Figure S4.** Alignment of *C. elegans* TRAP-1 and human TRAPα amino acid sequences.

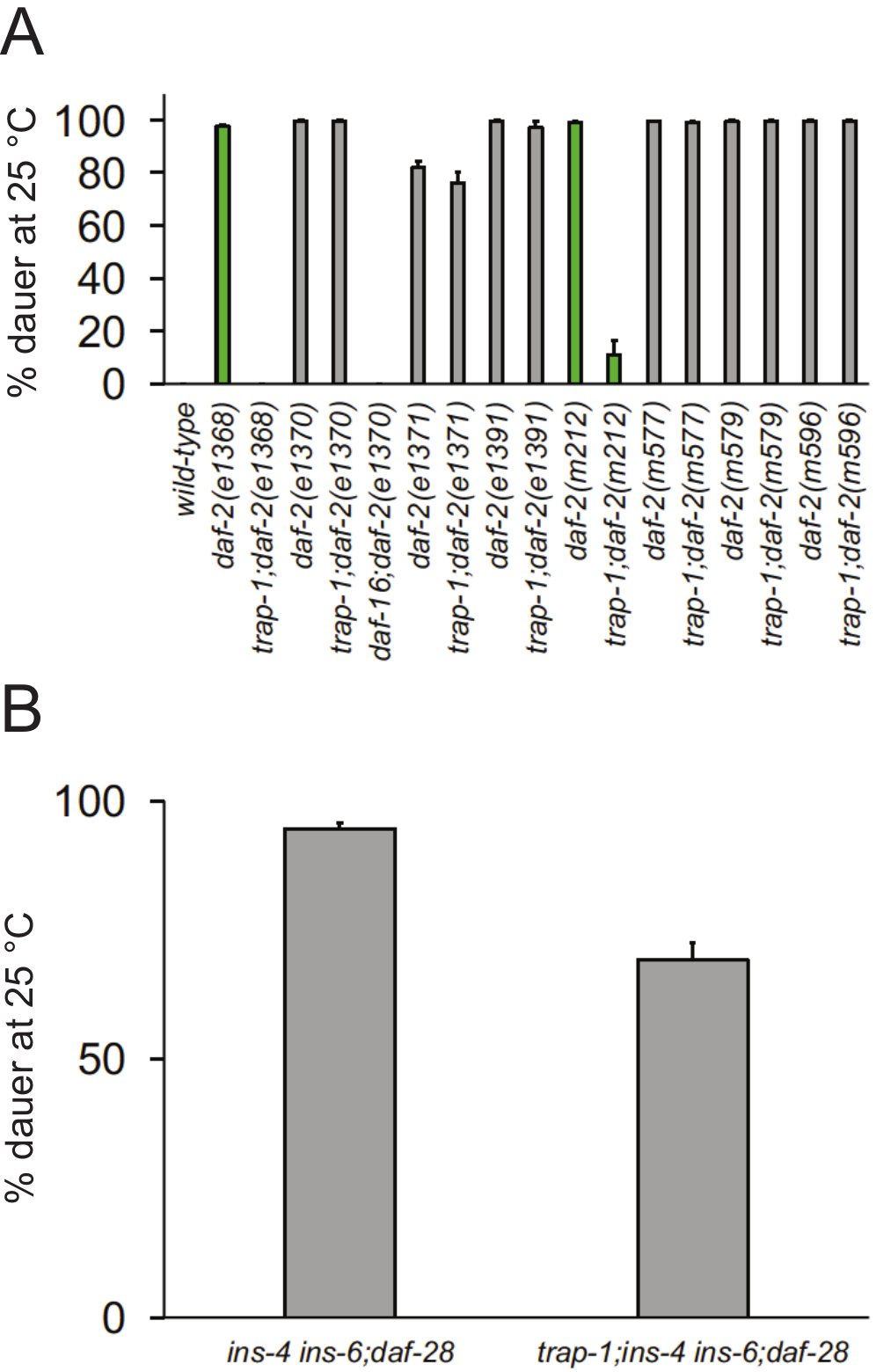

**Figure S5.** Effect of *trap-1* null mutation on dauer-constitutive phenotypes of A. eight *daf-2/InsR* loss-of-function alleles, and B. a triple mutant lacking the activity of the agonist insulin-like peptides INS-4, INS-6, and DAF-28 (p = 1.1e^-07^ by Student’s t-test). Error bars: s.e.m.

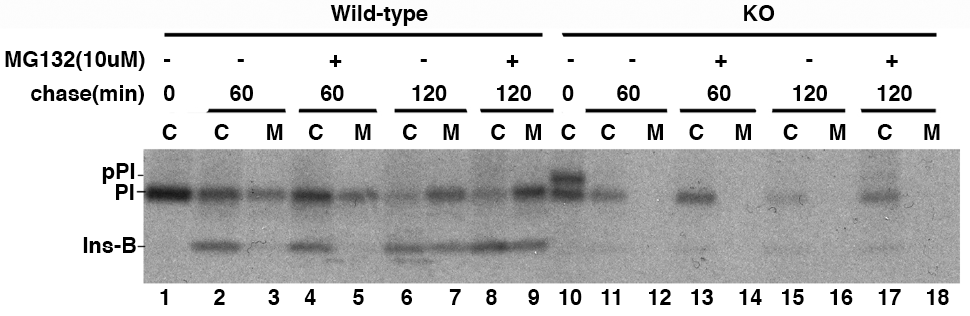

**Figure S6.** Turnover of newly synthesized proinsulin in TRAPα KO cells is partially dependent upon the proteasome. SDS-PAGE of anti-insulin immunoprecipitates of cell lysates (C) or conditioned media (M) from INS 832/13 wild-type or TRAPα KO cells was performed after pulse-labeling with ^35^S-Met/Cys and chase for the indicated times in the presence or absence of the proteasome inhibitor MG132. Abbreviations: pPI, preproinsulin; PI, proinsulin; Ins-B, insulin.

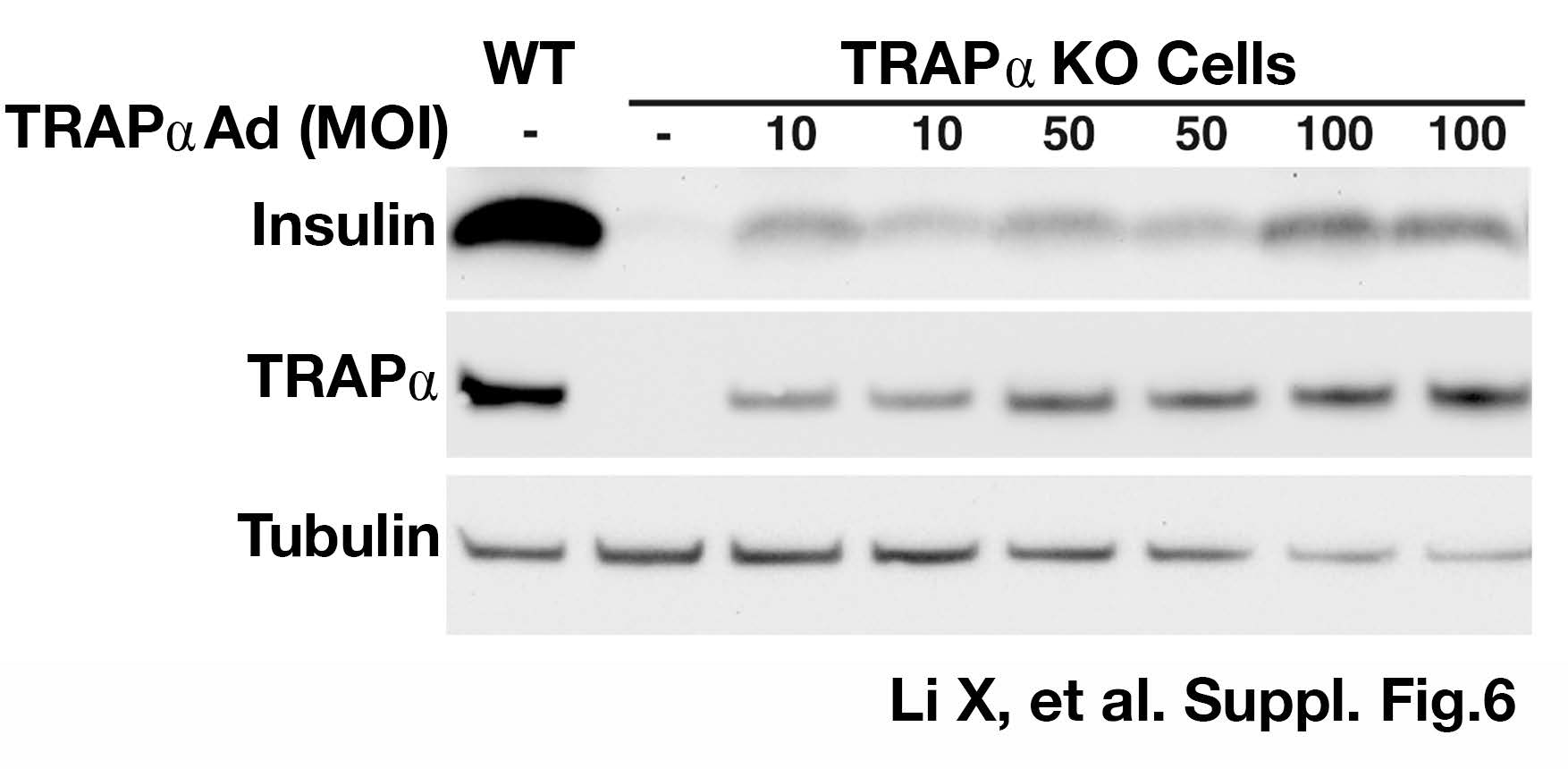

**Figure S7.** Re-expression of TRAPα in TRAPα KO cells rescues insulin production in a dose-dependent manner. TRAPα KO cells were infected with adenovirus expressing human TRAPα at increasing multiplicity of infection (MOI) as indicated. 48 hours after infection, cell lysates were subjected to SDS-PAGE and immunoblotted with anti-insulin, anti-TRAPα, and anti-tubulin antibodies. Duplicate samples of adenovirus-infected cells were analyzed.

| *daf-2/InsR* allele (Class) | Mutation | Human INSR residue/mutation | Location  (domain) | Disease  Association | Effect of mutation | Suppressed by *trap-1(null)*? |
| --- | --- | --- | --- | --- | --- | --- |
| *e1368* (1) | S573L | Not conserved | L2 | None | Not known | Yes |
| *e1371* (1) | G803E | G650 (none) | 2^nd^ FNIII | None | Not known | No |
| *m212* (1) | C883Y | C709 (none) | αCT | None | Not known | Yes |
| *m577* (1) | C1045Y | C825S | 2^nd^ FNIII | None | Not processed | No |
| *e1370* (2) | P1465S | P1236 (none) | kinase | None | Not known | No |
| *e1391* (2) | P1434L | P1205L | kinase | IRAN Type A | Decreased phosphorylation | No |
| *m579* (2) | R437C | R279C | CR | IRAN Type A | Reduced affinity for insulin | No |
| *m596* (2) | G547S | G393R | L2 | IR/LEPRCH | 90% decrease in cell surface receptors | No |

**Table S1.** Description of eight *daf-2/InsR* alleles assayed in Figure S5A. *daf-2/InsR* alleles have been grouped into two phenotypic classes, with Class 1 alleles exhibiting weaker mutant phenotypes than Class 2 alleles^40^. See text for details. Abbreviations: L2, second L domain; FNIII, fibronectin Type III; αCT, alpha-subunit C-terminal; CR, cysteine-rich; IRAN Type A, insulin-resistant diabetes mellitus with acanthosis nigricans Type A; IR/LEPRCH, insulin resistance/leprechaunism.
